# Whole-body gene expression atlas of an adult metazoan

**DOI:** 10.1101/2022.11.06.515345

**Authors:** Abbas Ghaddar, Erick Armingol, Chau Huynh, Louis Gevirtzman, Nathan E. Lewis, Robert Waterston, Eyleen J. O’Rourke

**Affiliations:** Department of Biology, College of Arts and Sciences, University of Virginia, Charlottesville, VA 22903, USA; Bioinformatics and Systems Biology Graduate Program, University of California, San Diego, La Jolla, CA 92093, USA; Department of Pediatrics, University of California, San Diego, La Jolla, CA 92093, USA; Department of Genome Sciences, University of Washington, Seattle, WA, USA; Department of Bioengineering, University of California, San Diego, La Jolla, CA 92093, USA; University of Virginia School of Medicine, Department of Cell Biology, Charlottesville, VA 22903, United States; Robert M. Berne Cardiovascular Research Center, School of Medicine, University of Virginia, Charlottesville, VA 22903, USA

## Abstract

Animals are integrated organ systems composed of interacting cells whose structure and function are in turn defined by their active genes. Understanding what distinguishes physiological and disease states therefore requires systemic knowledge of the gene activities that define the distinct cells that make up an animal. Towards this goal, this study reports the first single-cell resolution transcriptional atlas of a fertile multicellular organism: *Caenorhabditis elegans*. The scRNA-Seq compendium of wild-type young adult *C. elegans* comprises 159 distinct cell types with 18,033 genes expressed across cell types. Fewer than 300 of these genes are housekeeping genes as evidenced by their consistent expression across cell types and conditions, and by their basic and essential functions; 170 of these housekeeping genes are conserved across phyla. The 362 transcription factors with available ChIP-Seq data are linked to patterns of gene expression of different cell types. To identify potential interactions between cell types, we used the *in silico* tool cell2cell to predict molecular patterns reflecting both known and uncharacterized intercellular interactions across the *C. elegans* body. Finally, we present WormSeq (wormseq.org), a web interface that, among other functions, enables users to query gene expression across cell types, identify cell-type specific and potential housekeeping genes, analyze candidate ligand-receptors mediating communication between cells, and study promiscuous and cell-specific transcription factors. The datasets, analyses, and tools presented here will enable the generation of testable hypotheses about the cell and organ-specific function of genes in diverse biological contexts.

## Introduction

A wide range of different cell types sustains growth and reproduction in multicellular organisms. Even a simple animal like *Caenorhabditis elegans* develops according to a selected plan, survives biotic and abiotic stressors, discerns food quality, finds mates, escapes predators, and learns to associate environmental cues. In *C. elegans*, these functions are carried out by around 20 broadly-defined cell types and more than 150 specific cell types^1–3^. Underlying the morphological and functional differences between cells are cell-type specific networks of active genes. Thus, unveiling the molecular mechanisms underlying the functioning of multicellular organisms in physiological and pathological conditions requires a single-cell resolution catalog of the expressed genes and in turn, the genetic networks that define and are critical to maintaining cell identity and function. Such a catalog will facilitate future studies to assess how perturbations (genetic, chemical, or environmental) alter the genetic networks of cells and how these changes in turn result in phenotypes at higher levels of organization.

Recent advances in single-cell transcriptomics and *C. elegans* cell-dissociation protocols ^4^ have led to single-cell gene expression profiles of *C. elegans* embryos and larvae ^1,3–6^. Here we present the expression profiles of 159 cell types identified in wild-type and fertile *C. elegans* young adults. The single-cell resolution transcriptional map of the *C. elegans* adult adds several cell types present in the adult worms that are absent in the embryonic and larval stages, including various germ cells and cells involved in reproduction and egg laying.

We use this adult gene catalog to explore housekeeping genes, transcription factor (TF) associations with cell types, and cell-cell interactions. We identify genes that meet the canonical definition of a housekeeping gene, and are likely responsible for basic cellular maintenance across cell types and possibly kingdoms. We begin to elucidate, at a systems level, the relationship between the transcriptional programs and the identity of cells. We also infer cell-cell interactions between the cell types identified and predict the ligand-receptor pairs that promote these interactions. As a result, novel cell-type specific cell communication signatures are proposed, some of which we experimentally validate *in vivo*. Finally, we present a web interface, WormSeq, to mine our dataset, that together with the abundant literature and the genetic tools available to manipulate *C. elegans* will advance our understanding of the biology underlying the functioning of a multicellular organism and the perturbations that lead to its breakdown.

## Results

### Identification of over 163 distinct C. elegans cell types and subtypes

Wild-type hermaphrodite *C. elegans* were harvested as young adults (YA), as defined by vulva morphology and the presence of ≤5 eggs, dissociated into single cells (Fig. S1A), and subjected to scRNA-Seq using the 10X Chromium platform (See Materials and methods). Three independent biological replicates were collected and after the removal of low quality and damaged cells (see Materials and Methods), the dataset contained 154,251 cells that passed quality filters. The cells were then processed following the Monocle3 pipeline ^7^ and visualized using UMAP. After Louvain clustering the cells separated into 170 distinct clusters ranging from 21 to 5841 cells (Fig. 1A). Comparing replicates did not show batch-dependent differences in the average reads per cell (Fig. S1B) or genes per cell (Fig. S1C), and batch-dependent differences in the proportion of cells of different types were minimal (Fig. S1D), suggesting high reproducibility between independent experiments. Additionally, even prior to batch correction, cell type-specific gene expression profiles between biological replicates were highly correlated (Fig. S1E; Pearson correlation coefficient: 0.86-0.95), which suggests that although batch differences exist, well-controlled replicates accurately recapitulate average cell types and gene-expression profiles, and that the effect of cell type on the measured gene expression is stronger than the batch effect.

**Figure 1.**
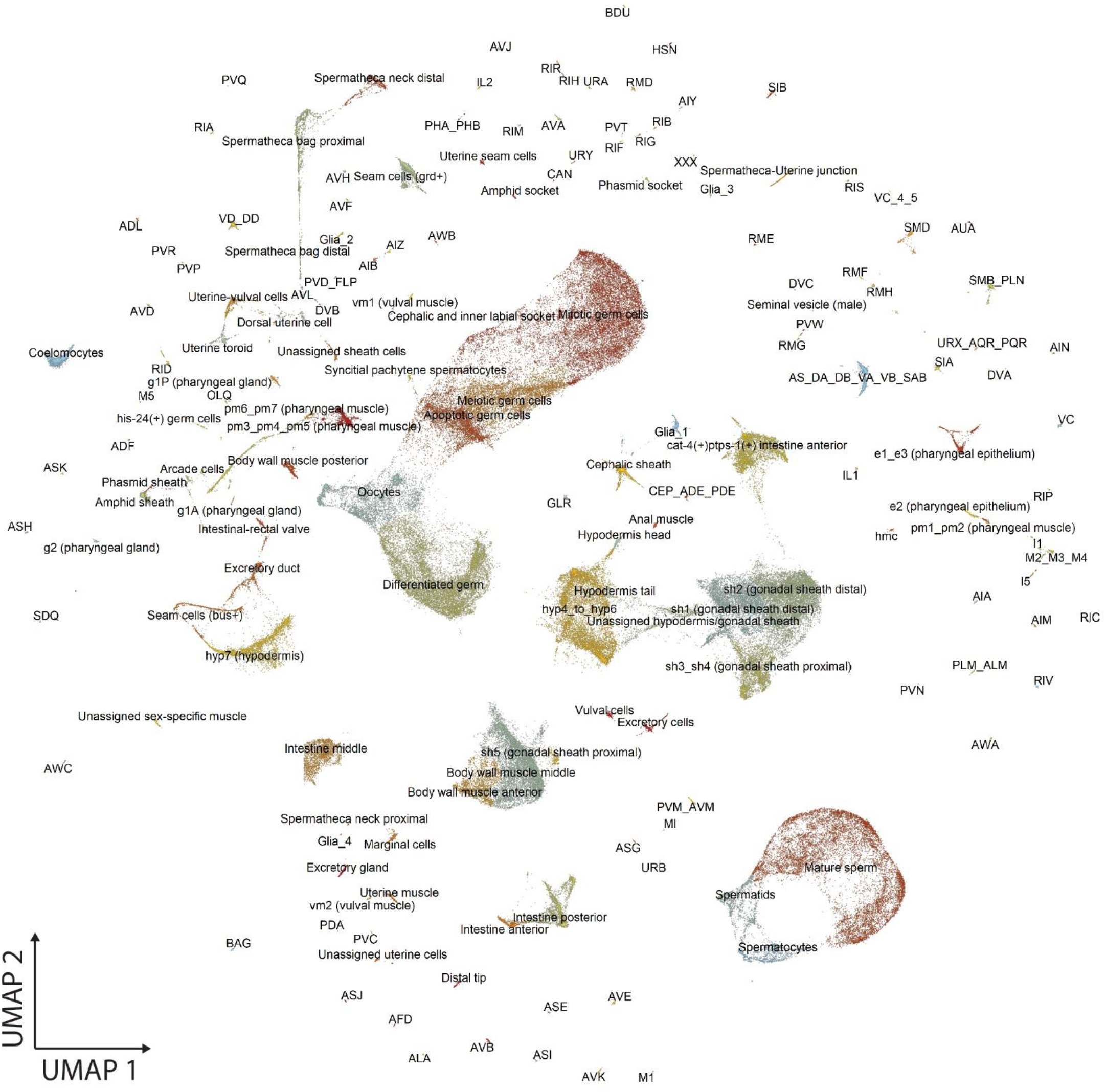
UMAP visualization of all identified cell types. UMAP reduction of 154,251 cells. Each dot represents a cell. Colors indicate distinct cell types and are used to facilitate distinguishing close clusters.

The clusters were annotated using a multi-pronged approach that took advantage of previously published scRNA-Seq data from *C. elegans* larvae ^1,3^ and the rich literature on *C. elegans* tissue and cell-specific markers ^8^. First, we generated a list of marker genes expressed in each cluster using Monocle3’s top_markers function (Table S1). We then searched for the broad cell types in which these marker genes were expressed in the CeNGEN app ^3^, as this dataset contains scRNA-Seq data from the L4 larval stage, the larval stage preceding the YA. This approach yielded broadly defined tissues. However, some clusters could not be confidently annotated using this approach because they lacked sufficient detail in the CeNGEN dataset (e.g. pharyngeal gland cells g1A vs g1P vs g2), and other clusters were not expected to be present in CeNGEN because they are adult specific (e.g. cells involved in egg laying). We therefore also manually identified gene markers using Wormbase. The gene markers used to annotate each cluster and the rationale behind every annotation can be found in Table S1.

We identified all expected broad cell types in the adult *C. elegans* hermaphrodite ^2^ including intestine, hypodermis, non-striated and body wall muscle, neurons, glial cells, pharynx, rectum and anus, seam, somatic gonad, vulva and uterus, excretory, coelomocytes, GLR cells, head mesodermal cells and XXX cells. More importantly, several major cell types could be further annotated into specific cell types as they had distinctive transcriptional profiles. The data was sufficiently exhaustive that we could identify cell types represented by a few or even one cell in the *C. elegans* adult. For example, we were able to identify specific pharyngeal gland cells (g1A, g1P and g2), individual gonadal sheath cells (sh1, sh2, sh3_sh4, sh5), and vulval muscle cells (vm1 and vm2). Finally, we identified 106 distinct neurons including 9 GABAergic, 47 cholinergic, 32 glutamatergic, 3 dopaminergic, 1 octopaminergic, and 2 serotonergic neuron types. Classifying the neurons by function, we found 30 sensory neurons, 20 motor neurons, 8 pharyngeal neurons, 34 interneurons, and 13 polymodal neurons. Overall, we defined 163 specific cell types (Table S1 & Fig. 1).

The analysis also revealed new sub-populations of cell types previously considered to be uniform. For example, we identified 5 clusters expressing distinct transcriptional programs belonging to the spermatheca. These distinctions were surprising given that the spermatheca is generally divided into three distinct components: the spermatheca neck, the spermatheca bag, and the spermathecal-uterine junction. Our results suggest that the spermatheca neck can be further subdivided into at least two populations of cells, named relative to the uterus: (i) spermatheca neck distal, which expresses *apx-1* and *let-502* ^9,10^, and (ii) spermatheca neck proximal, which does not express these markers. Similarly, the spermatheca bag can be subdivided into at least two subpopulations of cells: (i) spermatheca bag distal, which expresses *ajm-1* and *par-3* ^11^, and (ii) spermatheca bag proximal, which does not express these markers (further information about the rationale for this annotation can be found in Table S1). A single cluster corresponds to the spermathecal-uterine junction. Therefore, even in an organism with every cell anatomically mapped, potentially novel divisions of labor between cells can be uncovered using whole-body scRNA-Seq.

The initial list of cell types contained types not found in *C. elegans* adult hermaphrodites. One of these clusters expressed seminal-vesicle gene markers, which are exclusively found in males. We suspect these reflect the presence of rare males (≤0.01%) in our cultures ^12,13^. There were also cell clusters characteristic of the L4 stage (*e*.*g*., spermatocytes and spermatids), and of early embryos, which we postulate came from contaminating older adults in our starting populations. These observations make the total number of adult hermaphrodite cells 159, and indicate that droplet-based scRNA-Seq can capture underrepresented cell populations or subtle perturbations.

### Identification of housekeeping genes

Housekeeping genes can provide insight into intriguing biological questions including defining the genes under strongest selective pressure, or from a reductionist perspective, the genes essential to sustain life. Housekeeping genes also serve as references in various molecular and biochemical assays. However, it remains unclear whether commonly employed housekeeping genes, or any gene, meet the commonly used criteria to define housekeeping genes, namely consistent expression across cell types and conditions, essentiality, and conservation. To assess consistent expression, we employed two different criteria. We first applied a strict criterion: abundant expression within each cell type and expression across cells. For this, we created a gene-by-cell-type matrix to define for each cell type how many cells expressed a given gene (scaled TPM > 1). We then used density plots to visualize the prevalence of every gene across cells within cell types. A gene with a density plot skewed to the right (negative skewness score) is expressed in the majority of cells of most cell types (e.g., *ctc-3* in Fig. 2A). By contrast, a gene with a density plot skewed to the left (positive skewness score) is expressed in a minority of the cells of a few cell types (e.g., *sax-2* Fig. 2A). Only 53 genes had negative scores (Table S2) indicating that, in our dataset, very few genes meet the criteria of being ubiquitously and abundantly expressed across all cell types. However, this relatively small number of genes with negative skewness scores could be due to the technical limitations of scRNA-Seq (see Discussion). Nevertheless, these 53 genes were enriched in “basal cellular” functions including ribosomal activity, protein translation, and mitochondrial respiration (Fig. 2B), and in essential genes (lethal) as defined in partial ^14^ and full-genome RNAi screens ^15,16^ (Fig. 2C), two features in line with these genes being bona fide housekeeping genes.

**Figure 2.**
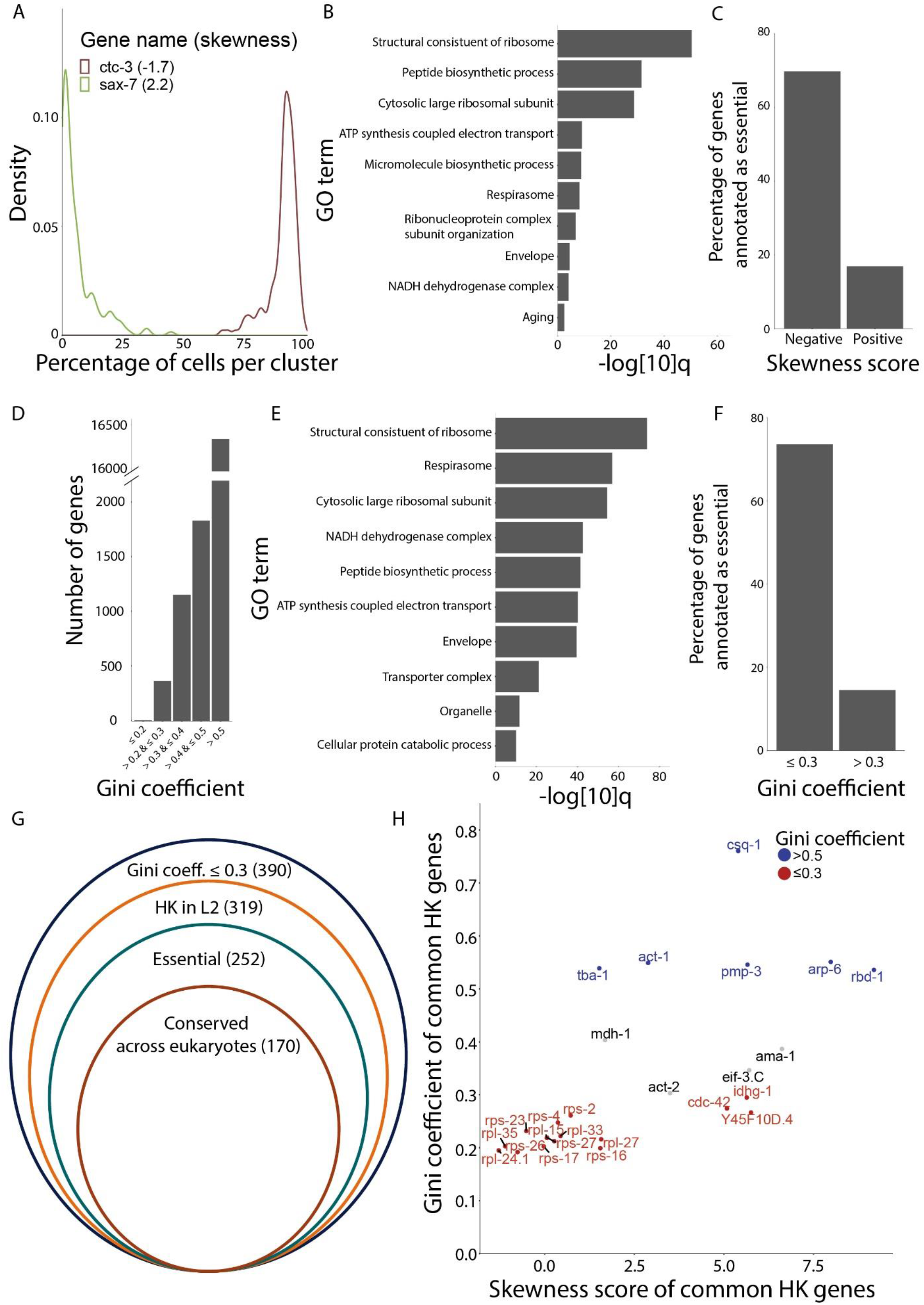
Identification of housekeeping genes using scRNA-Seq data. **(A)** Density plots of 3 genes illustrating the spectrum of skewness scores (in parenthesis), which represent relative abundance within and across cell types. **(B)** Gene Ontology (GO) enrichment analysis for genes with negative skewness scores (q-value threshold = 0.1). **(C)** Proportion of essential genes, defined as genes that are lethal when knocked down by RNAi, among genes with negative and positive skewness scores. Fisher’s exact test for enrichment p-value <2.2e-16. Note: this is the smallest p-value possible for this test. **(D)** Number of genes within each Gini coefficient (Gc) bracket: ≤ 0.2 perfect, >0.2 to ≤ 0.3 good, and >0.3 to ≤ 0.4 adequate expression equality. By contrast, Gc >0.4 to ≤ 0.5 indicates a big expression gap and >0.5 indicates a severe expression gap. **(E)** GO enrichment analysis for genes with low Gini coefficients (<0.3) (q-value threshold = 0.1). **(F)** Proportion of essential genes among genes with low (<0.3) or higher Gini coefficients (>0.3). Fisher’s exact test for enrichment p-value <2.2e-16. **(G)** Number of genes in our scRNA-Seq dataset with low Gc (<0.3), that showed housekeeping properties in the L2 worm13, were experimentally shown to be essential, and are conserved across species (see also Table S2). (H) Gini coefficient and skewness score of a set of commonly used housekeeping genes. Red font indicates genes with perfect or good expression equality (Gc ≤ 0.3) and blue indicates genes with a severe expression gap (Gc > 0.5). HK = housekeeping.

However, a housekeeping gene may not necessarily be abundantly expressed. An alternative metric of “housekeeping-ness” that would allow for this possibility, would be consistency of expression across cell types independent of abundance. To identify genes consistently, but not necessarily abundantly, expressed, we applied a metric of inequality called the Gini coefficient or Gc ^14^. Genes with lower Gc’s are expressed more equally across cell types, and genes with higher Gc’s are expressed in a more cell-type-specific manner. In our scRNA-Seq dataset, more than 90% of genes had Gc’s indicative of inconsistent expression across cell types (Gc ≥0.4). By contrast, 390 genes had Gc’s considered to represent good to perfect equality (<0.3) (Fig. 2D & Tables S2A and S2B), which suggests they might play a role in common or core cellular functions. In fact, the genes with low Gc (<0.3) were enriched in “basal cellular” functions, including ribosomal activity and mitochondrial respiration (Fig. 2E). Additionally, housekeeping genes are expected to be similarly expressed across conditions. As scRNA-Seq is not yet available for *C. elegans* subject to perturbations (genetic, chemical, or other), we used time, in the form of a different life stage, as the alternative condition. Specifically, we looked at the overlap between genes consistently expressed (Gc <0.3) in young adults and a list of putative housekeeping genes reported for the *C. elegans* L2 larvae ^14^. We found that all genes with high expression equality (Gs ≤0.2) and the majority of genes (319/390) with good expression equality (Gs <0.3) in the young adults were also consistently expressed across cell types in the L2 larvae (Table S2). Furthermore, the majority of these genes (252/319) were experimentally shown to be essential (lethal) in partial ^14^ and full-genome RNAi screens ^15,16^ (Fig. 2G). Finally, based on the conservation criteria defined in Tabach et al ^17^, we found that the majority of the low Gc genes (170/252) are conserved across animals, plants, and fungi (Fig. 2G; Table S2). Therefore, the 170-gene list meets several of the ascribed, but rarely tested, criteria that define housekeeping genes: (i) expressed consistently across cell types, (ii) expressed consistently across conditions (e.g. developmental stages), (iii) involved in basic cellular functions, (iv) essential for life, and (iv) conserved across species.

Next, we assessed the “housekeeping-ness” of 26 genes broadly used as “housekeeping” genes for normalization of gene expression in *C. elegans* ^18,19^. The majority of these genes (16/26) had Gc ≤0.3, indicating that these may be appropriate reference genes (Fig. 2H). In particular, 5 genes (*rpl-24*.*1, rpl-35, rps-26, rps-23* and *rps-17*) out of the 16 had a negative skewness score in our data, indicating that these 5 genes are not only consistently but also abundantly expressed across cell types (Fig. 2H). On the other hand, 6 genes (*rbd-1, tba-1, pmp-3, act-1, arp-6* and *csq-1*) had a Gc of more than 0.5; this severe expression gap indicates that these genes are inadequate normalization factors (Fig. 2H). In fact, all 5 genes are expressed in a tissue-specific manner (Fig. S2A-F).

### Inferring transcriptional regulators underlying cell identity

In line with the reported L2 single-cell data ^5^, we hypothesized that correlating the transcription factor (TF) binding patterns, as reflected in ChIP-Seq data ^20–22^, with gene expression profiles could give insights into the regulatory programs driving the gene expression profiles of the different cell types. For each of the 163 different cell types we constructed regression models to predict each gene’s expression as a function of the strength of the ChIP-Seq peak(s) proximal to its promoter region. After restricting correlations to TFs that were detectably expressed in our scRNA-Seq dataset (scaled TPM > 0), we ended up with 6691 TF binding-cell type expression associations (correlation > 0) (Fig. 3A; Table S3). To assess the validity of these associations, we tested whether the inferred TF association was able to predict cellular identity. We first clustered cell types using the TF scores alone (Fig. 3B). Then, we compared the resulting ‘TF association-based dendrogram’ to the ‘all expressed genes-based dendrogram’ using the entanglement score from the R package dedextend ^23^. Comparing the TF-based to the gene expression-based dendrogram yielded a low entanglement score (0.13), indicating high similarity (a score of <0.2 indicates high similarity). This high similarity means that TF activity largely drives cellular identity and, as such, it can predict cell-ontology relationships between cell populations.

**Figure 3.**
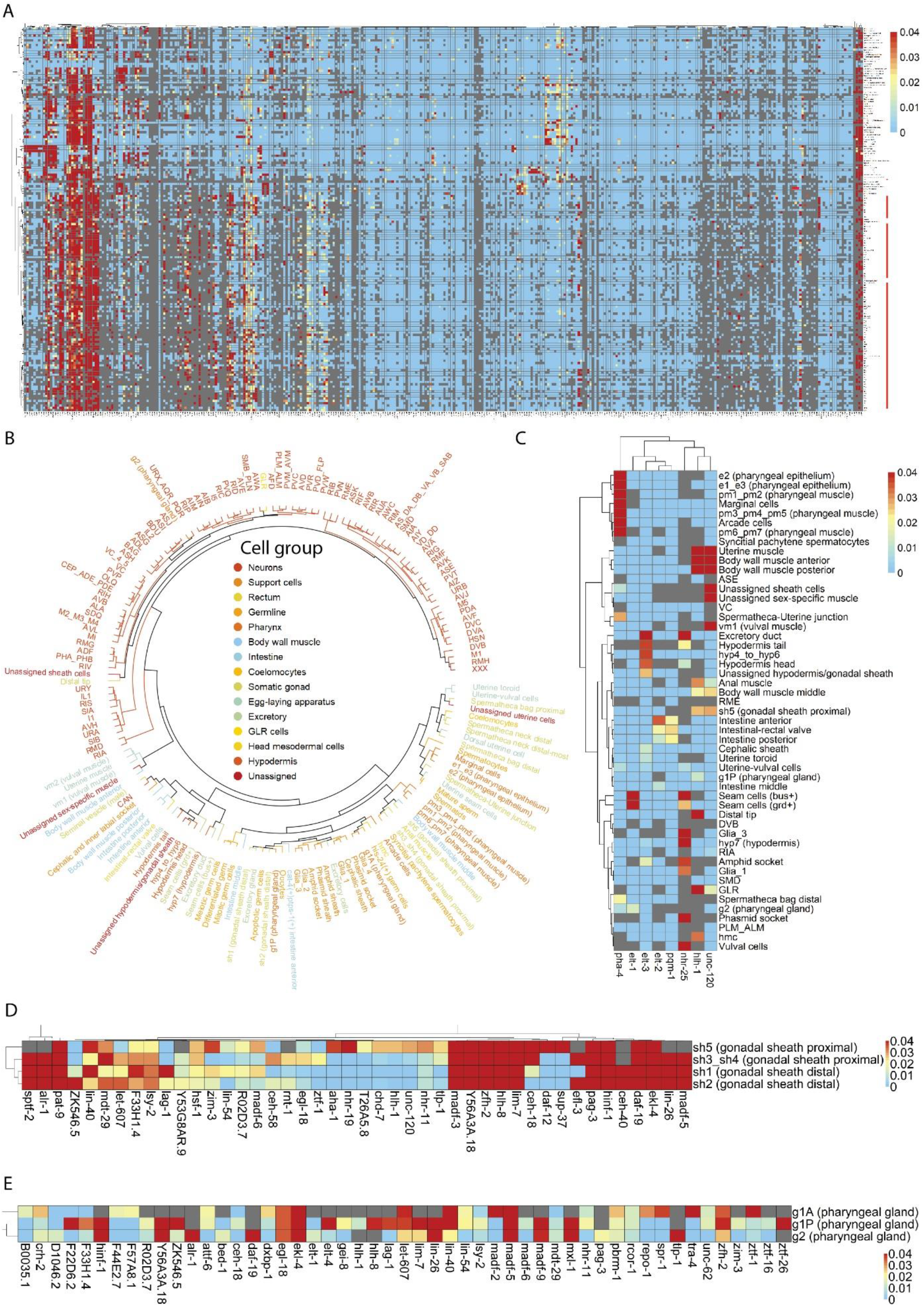
Inferring the transcriptional regulators of cellular identity. **(A)** Heatmap of the coefficients for TF_ cell type associations, with TFs on the x-axis and cell types on the y-axis. Hierarchical clustering of cell types and TFs was performed using the Ward.D2 method ^32^. The neurons (red label) cluster together in the bottom half of the map away from the other cell types. **(B)** A circularized dendrogram depicting the relationship between the cell types constructed using TF activity alone. Subtypes of cells are colored by broadly defined cell types. **(C)** Sub-heatmap showing that the analysis recapitulates previously known TF-cell type associations. **(D)** Sub-heatmap showing the predicted TF activity for all three pharyngeal gland subtypes. Only transcription factors that positively correlate with at least one of the cell types are shown here. **(E)** Sub-heatmap showing the predicted TF activity for all gonadal sheath subtypes. Only transcription factors that positively correlate with at least one of the cell types are shown here.

The regression analysis was able to recapitulate known cell type-TF associations (Fig. 3C) including body wall muscle with *hlh-1* and *unc-120* ^24^, hypodermis with *nhr-25* and *elt-3* ^25,26^, seam cell with *elt-1* ^27^, intestine with *elt-2* and *pqm-1* ^28,29^, and pharynx with *pha-4* ^30^. The regression analysis also suggested several previously unreported regulatory relationships (Fig. 3C). For instance, *nhr-25* appears to be specifically active in the hyp7 hypodermal cells, while elt-3 is predicted to be active in hyp4_hyp6 head and tail hypodermal cells but not in hyp7. We also found that *unc-120* shows a high regression coefficient in several muscle cells including all sex-specific muscle cells as well as body wall muscle while *hlh-1* had a high regression coefficient in anal and uterine muscle cells in addition to body wall muscles. We were also able to identify common and distinctive TFs between closely related cell subtypes. For example, our analysis predicts that the TFs *madf-5, ekl-4*, and *egl-18* are common to all three pharyngeal gland subtypes while *ztf-1* is specific to g1A, *daf-19* is specific to g2, and *ztf-19* is specific to g1P cells (Fig. 3D). Similarly, the TFs *hinf-1* and *pat-9* are common to all gonadal sheath cells (sh1, sh2, sh3_sh4 and sh5). By contrast, *lag-1* and ZK546.5 are specific to sh1 and sh2; *ztf-1* is specific to sh3_sh4; and *daf-12* and *sup-37* are specific to sh5 cells (Fig. 3E). While additional experiments will be needed to validate these inferred relationships, the results highlight the potential of combining expression data with TF binding data to advance our understanding of the transcriptional programs responsible for maintaining the identity of even closely related cells.

Finally, we looked into the relationship between these predicted TF associations and cellular identity. Specifically, we tested whether combining our scRNA-Seq data, ChIP-Seq data, and *in vivo* experimental data enabled the identification of the molecular targets through which a TF of interest maintains the functional identity of a cell. For instance, our data predict that the TF *dsc-1* is active in the anal muscle. In agreement with this association, RNAi against *dsc-1* causes constipation and shorter defecation cycles ^31^. Correspondingly, we identified 1089 DSC-1 gene targets expressed in the anal muscle. Among the 1089 genes, 29 were known to contribute to normal defecation in *C. elegans*, a significant enrichment as measured by a Fisher’s exact test (p-value = 3.183e-09, Table S4). From the remaining DSC-1 targets, at least 70 genes are known to be required for muscle activity ^8^, and hence, may similarly contribute to defecation. These are now testable hypotheses that experts in the field can pursue to define anal muscle function through the activation of a transcriptional program that is, at least partly, cell-autonomously executed.

### Whole-body reconstruction of cell-to-cell interactions

Cell-cell interactions (CCIs) are critical to the maintenance of the tissues and organ systems that sustain life. We previously developed the tool *cell2cell* to infer CCIs from the expression of ligand and receptor encoding genes across cells in single-cell transcriptomics datasets, and curated a database of ligand-receptor pairs to study CCIs in the *C. elegans* L2 larvae ^33^. Here, we use these computational resources, together with a permutation analysis ^34,35^, to identify CCIs between all 163 cell types and subtypes across the whole-body of young adult *C. elegans* and to predict the likely molecular drivers of those interactions. A large matrix of putative interactions was obtained (see the WormSeq to browse these interactions), below we discuss a few illustrative examples.

*Cell2cell* was able to identify previously validated CCIs. For example, *cell2cell* predicts that the distal tip cells interact with germ cells ^36^, and that a major driver of this CCI is the molecular interaction between the ligand *lag-2* and the receptor *glp-1. Cell2cell* also predicted several unreported molecular interactions between cell types. For example, the ligand-receptor pair *nlg-1*/*nrx-1* is the highest-scored driver of the interaction between AVA and various motor neurons. Although AVA neurons are involved in touch-induced locomotion ^37^ and *nlg-1* RNAi treated worms are resistant to touch-induced locomotion ^38^, it was not known which signaling molecules produced by AVA would contribute to the touch response. However, the combination of the published data with the *cell2cell* results led us to hypothesize that the interaction between the AVA-generated NLG-1 ligand and the NRX-1 receptor in motor neurons contributes to the touch response. Similarly, *cell2cell* predicts that *sax-7*/*pat-2* and *sax-7*/*pat-3* contribute to the interaction between DVB neurons and anal muscle cells. In support of these molecular interactions, DVB neurons innervate the anal muscle to regulate defecation. Furthermore, *sax-7* mutant worms have reduced defecation rates relative to wild-type worms ^39^. However, it was not known the site of action of *sax-7* as it relates to the control of defecation, or which receptor would perceive this signal in the anal muscle. Together, the *cell2cell* analysis and the published work enable us to hypothesize that the molecular interaction between DVB-generated SAX-7 and PAT-2/3 in the anal muscle contributes to normal defecation in *C. elegans*. In addition, *cell2cell* predicts that *sax-7/pat-2* and *sax-7/pat-3* mediate a functional interaction between VC4 and VC5 neurons and the sex-specific muscles. This prediction is supported by the fact that *sax-7* mutants are also egg-laying defective ^39^. These results suggest that the expression of the *sax-7* ligand in VC4 and VC5 is necessary for normal egg laying. These examples illustrate the power of *cell2cell* in combination with permutation analysis to predict the molecular drivers of biologically relevant cell-cell interactions.

Nevertheless, although *cell2cell* can make meaningful predictions of individual LR pairs, intercellular interactions are often driven by simultaneous interaction between multiple signals and receptors, some of which could counteract each other or change signs while carrying out different biological functions. To account for these and other complexities, we next used *Tensor-cell2cell*, an unsupervised machine-learning method that identifies patterns of cell-cell communication, and reports them as factors or signatures that summarize the operative LR pairs and cell types involved in an associated biological process ^40^. Using our whole-body scRNA-Seq data, we identified 11 unique signatures, each capturing a combination of ligand-receptor pairs and groups of cell types carrying out a biological function (Fig. 4A-C, Table S5). Validating the *Tensor-cell2cell* approach, some of the identified signatures were well-documented in the literature. For instance, Signature 10 (Fig. 4A-C) predicts a functional interaction between several neurons and germ and intestinal cells through the insulin-signaling pathway. On the ligand side, the analysis predicts that insulin-like peptides (ILPs) are mainly produced by neurons (Fig. 4A). In support of this Tensor-cell2cell prediction, multiple labs have shown that, with a few exceptions, insulin-like peptides (ILPs) are generated by neurons ^41–45^. However, our analysis goes further by predicting the following neuronal subtypes as the main generators of ILPs in fed young-adult *C. elegans*: ADF, AFD, AIA, AIN, ASE, ASI, ASJ, ASK, AWA, AWB, AWC, RIR, URB, and URX_AQR_PQR (Fig. 4A). Additionally, weaker ILP production is predicted to occur in ADL, ASG, ASH, BAG, M1 and RMH neurons. Furthermore, we can assign specific ILPs to specific neurons. For example, *ins-28* is prominently expressed in AIA, AIN, AWA and M1 neurons, while *ins-6* is more prominently expressed in ASI and ASJ neurons (Fig. S4A). On the receptor side, the most prominent receiver (receptor-producing) cells enriched in this interaction are intestinal and various germ cells (Fig. 4B). Correspondingly, several groups demonstrated the presence and functional relevance of the sole *C. elegans* insulin receptor, *daf-2*, in the germline ^46^ and intestinal cells ^47^. Additionally, we find several neurons enriched as insulin-signaling receiver cells (Fig. 4B), which is supported by published work showing that insulin-signaling also mediates interneuronal communication ^41^. For example, we recapitulated the insulin-signaling mediated interaction between AIA neurons and ASE neurons, which is required for salt chemotaxis learning ^48^. Specifically, AIA neurons prominently express ins-1 and ASE neurons express *daf-2* (Fig. S4E & F), which are both required for salt chemotaxis learning as previously reported ^48^. The results also predict novel insulin-signaling communication between the various ILP-producing neurons and a few receptor-producing neuronal subtypes including the *daf-2*/*daf-4* expressing neurons summarized in Table S4.

**Fig. 4.**
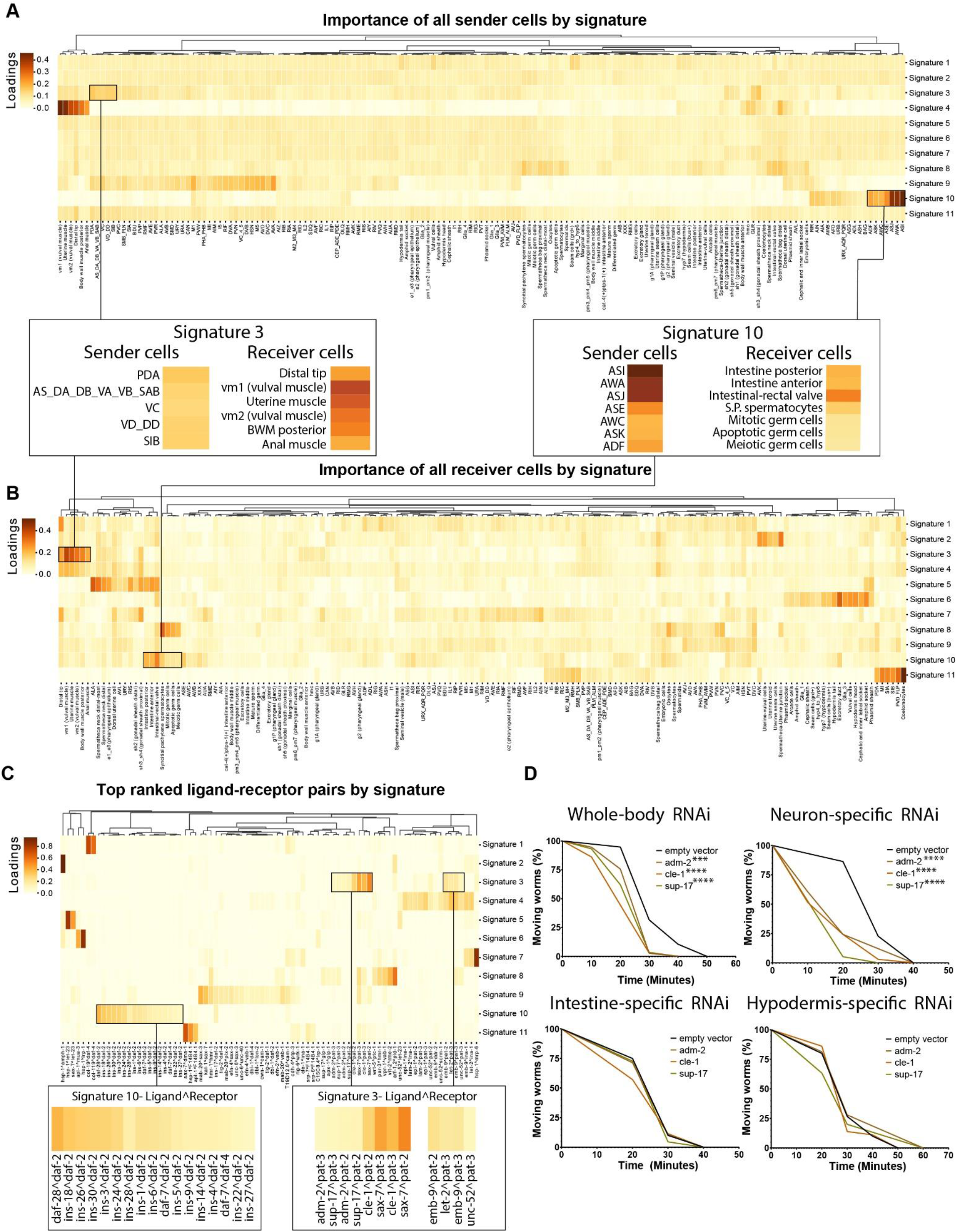
Identification of cell-type specific communication signatures using Tensor-cell2cell. **(A)** The heatmap shows which cell types play a major role as sender cells per signature. Inset shows which cell types are important as senders for signatures 3 and 10. **(B)** The heatmap shows which cell types are identified as main receiver cells per signature. Inset shows which cell types are important as receivers for signatures 3 and 10. **(C)** Heatmap showing which ligand-receptor pairs are the main mediators per signature. Inset shows which ligand-receptor pairs are important for signatures 3 and 10. In panels **(A-C)**, loadings represent the importance that *Tensor-cell2cell* assigned to each element within their respective signature. Panel **(C)** only shows ligand-receptor pairs that are important in at least one signature (loading value > 0.1). **(D)** Curves depict the time it takes for RNAi-treated worms to become paralyzed after levamisole treatment. *** p-value < 0.01; *** p-value < 0.001

*Tensor-cell2cell* was also able to predict novel cell communications signatures. For example, Signature 3 predicts that the ligands *adm-2, cle-1, emb-9, let-2, sax-7, sup-17*, and *unc-52* and their corresponding receptors *pat-2* and *pat-3*, mediate the interaction between motor neurons (senders) and muscle cells (receivers). Signature 3 specifically predicts that these ligand-receptor pairs mediate the interaction between the AS, DA, DB, DD, PDA, SIA, SIB, VC, and VD neurons and muscle cells from the body wall, anus, vulva, and uterus. Reassuringly, some of these predictions appear to be supported by experimental results. For example, CLE-1 is enriched in neuromuscular junctions ^49^, and *pat-2, pat-3, emb-9, let-2*, and *unc-52* contribute to muscle function ^50,51^. However, the site of expression and neuro-muscular function of the *adm-2, cle-1*, and *sup-17* ligands has not been reported. We, therefore, used a levamisole-sensitivity assay to test the *Tensor-cell2cell* prediction that inactivation of these three ligands may affect neuromuscular junction function. Levamisole is an acetylcholine receptor agonist that causes continued neuronal stimulation of the muscles, leading to paralysis ^52,53^. Resistance or hypersensitivity to levamisole is indicative of a neuromuscular junction dysfunction ^54^. We performed whole-body and tissue-specific RNAi knockdown of the ligands of interest starting at the young L4 stage, and when they reached the young adult stage we treated them with levamisole. As predicted by Signature 3, intestine- and hypodermis-specific ligand knockdown did not alter sensitivity to levamisole. By contrast, whole-body and neuron-specific knockdown of all three tested ligands resulted in levamisole hypersensitivity (Fig. 4D). Interestingly, the paralysis phenotype was more pronounced in a worm strain (TU3401; Fig. 4D) engineered to promote RNAi knockdown specifically in the neurons of *C. elegans* ^55^. Importantly, animals treated with RNAi against the ligands showed normal chemotaxis to sodium salts, which is another reported function of the neurons expressing these ligands (PDA,VC, VD DD, SIB) (data not shown), implying that knocking down the tested ligands does not cause pleiotropic dysfunction of the relevant neurons. Altogether, the match between the *Tensor-cell2cell* results and functional analyses suggests that this analysis is capable of generating meaningful hypotheses about the molecules driving cell-to-cell and cell-to-cell-to-function interactions in *C. elegans*.

### WormSeq app: Explore the whole-body transcriptional landscape of the adult C. elegans

To make the scRNA-Seq data more accessible to non-coding users, we created an RShiny app we called WormSeq. This resource is available as a web application and can be accessed using this link: wormseq.org. WormSeq has several features, including single-cell dot plots, and heatmap visualization of gene expression by count and by percentage of cells expressing a gene. Users can identify cell type specific gene markers by browsing gene marker tables or by using percentage gene expression per cell type. The interface also enables the identification of genes expressed in one cell type but not another one. Users interested in the abundance or consistency of gene expression can also browse genes by skewness score or Gini coefficient. The app also allows users to browse the regulatory program analysis data and identify transcription factors enriched in cell types of interest. Finally, users can also browse the *cell2cell* analysis and identify the list of interactors driving communication between two cell types of interest.

We anticipate that this tool will enable the generation of numerous testable hypotheses across fields of study.

## Discussion

We present here a comprehensive single-cell atlas of a wild-type adult *C. elegans*. Although single-cell transcriptional atlases have been generated for other metazoans, including mice and humans ^56,57^, this scRNA-Seq dataset is unique due to the following: (i) it derives from three independent populations each composed ∼100,000 animals, (ii) it was obtained from hermaphroditic, genetically homogeneous animals, which entails lower expression noise than what can be achieved in gonochoric species, and (iii) the soma, tissues, and organs of each and all adult *C. elegans* have the same cell types and number of cells (*e*.*g*., 95 body muscle cells in total). Such redundancy yielded a high-resolution scRNA-Seq dataset that captures all cell transcriptomes including those underrepresented in the starting worm populations (*e*.*g*., male cell types). Additionally, although current scRNA-seq protocols capture only a small fraction of the total RNA molecules per cell, our oversampling of *C. elegans* cells (total of 154,251 cells) and the aggregation of cells from the same cell type, enabled the reconstruction of a representative transcriptional profile for each cell type composed of a median number of 9,626 genes per cell type (scaled TPM ≥ 1). This level of gene activity per cell type poses interesting questions for future investigation. Which of these genes are required to maintain cell identity and function? Which ones are part of transcript reservoirs ready to act upon stress or other contexts? Which ones reflect biological or experimental noise? A more general limitation of any RNA analysis is that mRNA expression may or may not reflect protein abundance. In fact, previous studies have shown that the correlation between mRNA levels and protein levels can be poor ^58–60^. Therefore, incorporating proteomic data, and in the future single-cell proteomic data, is anticipated to increase the accuracy of functional predictions.

Despite the technical limitations, the *C. elegans* transcriptional atlas reported here is composed of the gene expression profiles of 163 distinct cell types. In addition to identifying most known *C. elegans* cell types, our annotation revealed differences in expression profiles between cell types previously assumed to be the same due to morphological similarity. For example, *apx-1* and *let-502* expressing in the *distal* but not in the *proximal spermatheca neck* together with the reports showing that whole-body RNAi against *apx-1* ^61^ and *let-502* ^62^ lead to dysregulation of the expansion of the germline, suggest that the *distal spermatheca neck* cells may contribute to tumorous processes in the germline. Therefore, the transcriptome-based annotation of *C. elegans* cells presented here opens doors to new cell-specific biology.

In this study, we also use the single cell data to begin to address three fundamental questions about the relationship between gene expression and cellular function: Which, if any, genes meet the definition of housekeeping gene? What transcriptional programs generate and maintain cellular identity in *C. elegans*? Which genes mediate the interactions between cells in this metazoan?

### What genes are housekeeping?

Using scRNA-Seq we were able to identify genes consistently expressed across cell types, and, hence, directly test for a feature commonly attributed to housekeeping genes. We scored all genes in our dataset based on the abundance and consistency of expression across cell types using a skewness score, or only for consistency of expression using a Gini coefficient. Although skewness score and Gini coefficient positively correlate with each other (Pearson correlation coefficient = 0.66), the Gini coefficient is more likely to have fewer false negatives since housekeeping genes are not necessarily abundantly expressed. Supporting the use of the more permissive Gini coefficient to identify housekeeping genes, the resulting list of consistently expressed genes (Gini coefficient ≤0.3) is enriched in genes essential to *C. elegans* survival (252/319) to an extent similar to the most restrictive skewness set (37/53). Furthermore, these genes are consistently expressed in two very distinct ontogenetic stages, the L2 larvae and the adult *C. elegan*s, showing that the Gini coefficient analysis applied to one condition can capture genes consistently expressed across conditions.

Because our housekeeping genes analysis is limited to the N2 wild type *C. elegans* strain, and it compares only two developmental stages (L2 larvae and young adult), it may be too inclusive. Additional scRNA-Seq experiments in other genetic backgrounds, developmental stages, or in the presence of abiotic or biotic stressors may further narrow the number of housekeeping genes, or even challenge the concept of housekeeping gene altogether. Nonetheless, the 252 genes found in the Gini analysis represent a solid beginning. As expected, these 252 genes are enriched in basic cellular functions including mitochondrial function, protein synthesis (*e*.*g*., ribosome and protein translation), and protein stability (*e*.*g*., chaperones). However, we did not find DNA synthesis/replication genes in this set, likely a reflection of the fact that apart from the germline, cells in the adult *C. elegans* are post-mitotic. We also did not find enrichment for RNA synthesis genes even though at least some of these genes are well represented across cell types (*e*.*g*., the RNApol encoding gene *ama-1*). Nevertheless, of the 252 genes consistently expressed in *C. elegans*, we found that 170 are conserved in organisms ranging from yeast to rice and humans (Table S2), and as such, they could be part of the core of genes indispensable to build and maintain a eukaryotic cell.

### What transcriptional programs generate and maintain cellular identity in C. elegans?

Our regression analysis of ChIP-Seq data with the cell type-specific gene expression profiles revealed 6691 TF-cell type associations, some of which were known, and as such, validate the approach. In line with the high resolution of our dataset, the TF-cell type association analysis identified TF associations unique to even closely related cell subtypes. In WormSeq, the web interface accompanying this study, users can search for all the TFs predicted to be active in each cell type, as well as all the cell types in which a given TF is predicted to be active. As the number of TFs with ChIP-seq data increases from the 362 TFs used here to encompass the 900 or so TFs predicted to exist in *C. elegans* ^63^, the association between TF binding patterns and cell-specific transcriptional profiles should become even more powerful in revealing regulatory relationships. Also, while we only evaluated the activating role of TFs, other advances may permit the investigation of negative regulators.

Through combining published ChIP-Seq, our scRNA-Seq data, and published functional data, we proposed cell type-TF-targets triads that may mediate cellular function and morphology. For example, knockdown of the gene encoding the TF *dsc-1* and of several of its ChIP-seq targets can alter defecation cycles in the worm ^31^. Our analysis predicts that *dsc-1* is active in anal muscle and that 36 of its downstream targets involved in defecation are expressed in the anal muscle. Therefore, we hypothesize that DSC-1, its 36 downstream effectors, and likely another set of 70 target genes that yield more general muscle phenotypes but are expressed in the anal muscle, orchestrate a cell-autonomous transcriptional program critical for normal defecation in *C. elegans*. The results presented in this study promise to accelerate advances by reducing the number of candidate genes for functional studies and by placing molecular players in their anatomical sites of action.

### Which genes mediate the interactions between cells in C. elegans?

We used the algorithm *cell2cell* and a curated list of ligand-receptor (LR) pairs to identify potential gene pairs mediating the interactions between the cells of the young-adult *C. elegans*. We also used a permutation analysis and *Tensor-cell2cell* ^40^ to identify cell-type specific communication signatures. Validating our approach, we detected known CCI-LR associations including insulin signaling mediating the communication between neurons and germline, and neurons and intestine. Additionally, even for this well-characterized communication pathway, our analyses provided new testable hypotheses including the specific neurons that produce insulin-like ligands.

Our predictions, however, are only based on the expression of LR pairs without accounting for spatial constraints. This omission may lead to predictions that do not match the biology, most notably membrane-bound LR pairs predicted to mediate the interaction between physically distant cells. For this reason, in our web interface WormSeq, we included a feature that allows users to browse our CCI-LR predictions by LR class: (i) membrane-bound, (ii) ECM component and/or (iii) secreted.

Overall, this study is a step forward toward identifying the key molecular players whose function or dysfunction defines the functional status of the intra and inter-cellular gene networks that constitute an animal. Putting together: (i) genetic tractability, (ii) stereotypical number and physical interactions between cells dissected to the level of synapses, (iii) known lineage for every cell in the body, and (iv) existing scRNA-Seq datasets of the wild-type embryo, two larval stages, and now the adult, will enable the development of tools that track gene expression across space and time for a whole living animal, as well as predictive models of cellular, tissue, organ, and ultimately whole-animal function. These tools can help many fields in at least two ways: (i) they may tell us how much information and of which kind is necessary to develop models that can accurately predict the effect of perturbations (biological, chemical, physical, or others), and (ii) the tools themselves may serve as the basis or guide the development of predictive tools for more complex organisms.

## Materials and Methods

### C. elegans strains and husbandry

*C. elegans* N2 (Bristol, UK), WM118 (rde-1(ne300);neIs9 [myo-3::HA::RDE-1 + rol-6(su1006)]), MGH171 (sid-1(qt9);alxIs9[vha-6p::sid-1::SL2::GFP]), JM43 (rde-1(ne219);xkIs99[wrt-2p::rde-1::unc-54 3’UTR]), TU3401 (sid-1(pk3321);uIs69 [pCFJ90 (myo-2p::mCherry) + unc-119p::sid-1] were obtained from the Caenorhabditis Genetics Center (CGC). All strains were typically grown at 20°C on NGM plates seeded with *E. coli* strain OP50. Bacterial strains used for RNAi were obtained from the Ahringer library ^64^.

### Sample preparation and scRNA-Seq

A synchronous population of L1 worms was obtained by double bleaching gravid N2 *C. elegans* with hypochlorite followed by 4 washes in S-buffer. The released eggs were then allowed to hatch in the absence of food in S-buffer over a period of 18 hours. Approximately 100,000 synchronized L1 worms were then grown in NGM plates seeded with HT115 bacteria at 20°C for approximately 55h. 55h post-seeding, worms were staged under a microscope to ensure that the bulk of the population had reached the young adult stage. Young adult worms were then harvested in S-buffer and then centrifuged at 1,300g for 1 min. The worm pellet was washed until the suspension was no longer turbid (2-3 times) and then transferred to a 1.5mL Eppendorf tube. The cuticle was then disrupted by incubating the worms in 200μL SDS-DTT (20mM HEPES pH8.0, 0.25% SDS 200 mM DTT, 3% sucrose) ^65^ for 4 minutes. Immediately after SDS-DTT treatment, 800 mL of egg buffer was added to the treated worms, the worms were centrifuged, the supernatant was aspirated and the worm pellet was washed 5 times in egg buffer (118 mM NaCl, 48 mM KCl, 2 mM CaCl2, 2 mM MgCl2, 25 mM HEPES, at osmolarity of 340 Osm). After the final wash, egg buffer was added to a final volume of 1mL and the worm solution was then transferred to a 15-mL conical tube. 500μL of 350 units/mL Pronase (EMD Millipore Corp) was added and the worms were then dissociated into single cells by passing them through a 21-gage needle about 20 times. The worm/cell lysate was centrifuged at 4°C for 1 min at 200g and then most of the supernatant, containing dissociated cells, was transferred to a new 15-mL conical tube leaving behind enough liquid for a second round of dissociation. After passing the worm lysate through the needle for a second time, the samples were centrifuged (4°C for 1 min at 200g) and then the supernatant was transferred to the same tube containing the cells from the first transfer. The cells were then centrifuged at 4°C for 5 min at 500g and the cell pellet was washed 3 times in egg buffer containing 1% BSA gently pipetting the cells with wide-end tips. Finally, to separate single cells from bigger chunks of tissue, the cell suspension was gently passed through a 10μm filter.

For single cell capture, 14,000 *C. elegans* cells were mixed with the reverse transcriptase solution and then loaded onto each channel of the 10x Chromium Controller. The libraries were then built following the Chromium Next GEM Single Cell Kits v3.1 published protocols and then sequenced on an Illumina NextSeq 500 platform.

### scRNA-Seq data processing

The scRNA-Seq data was first processed following the CellRanger pipeline. Reads were mapped to a modified version of the WormBase WS260 reference transcriptome that had transcript 3′ untranslated regions extended by 0 to 500 base pairs ^1^. To distinguish cells from empty droplets we used the knee plots reported by CellRanger to set a UMI threshold below which droplets were considered empty. The expression matrix generated by CellRanger was then decontaminated for ambient RNA using DecontX ^66^. We then followed the Monocle3 pipeline to perform dimensionality reduction and clustering ^7^. First, we combined all three biological replicates into a single cds object. We then used Monocle3’s preprocess_cds function (method = “PCA”, num_dim = 200) which normalizes the data by log factor and generates a lower dimensional space for downstream dimensionality reduction. Next, we used Monocle3’s align_cds function to perform further background correction and remove unwanted batch effects which we noticed came mostly from the different samples. We then performed UMAP dimensionality reduction on the matrix using Monocle3’s reduce_dimension function run with default parameters. Finally, we used Monocle3’s cluster_cells function to define individual clusters of cells using the Louvain algorithm (k=50).

After clustering, we noticed that there were clusters containing mostly cells with a high mitochondrial fraction (mitochondrial-only umi/total umi > 0.2). These clusters were removed since high mitochondrial fraction is an indication of damaged cells ^67^. We then re-performed dimensionality reduction and clustering on the remaining cells as described above. We also noted that some cells labeled “Intestine middle” prominently expressed hypodermal gene markers and were therefore removed from the data since they were likely intestine-hypodermis doublets.

### Cell type annotation

To annotate the different clusters of cells with their corresponding cell types we used Monocle3’s top_markers function to identify for every cluster a list of 10 gene markers. We then used the CeNGEN app ^3^ to broadly define where these genes are typically expressed in the L4 worm. In addition to the L4 data, we used gene markers identified through scRNA-Seq of L2 worms ^1^. The annotation of the L2 worms was more detailed than the CeNGEN data and allowed us to more carefully annotate several clusters. We also used Wormbase to identify gene markers for cell types that were absent from the L2 and L4 data and those that could not be confidently annotated using the L2 and L4 data alone. A detailed rationale for the annotation can be found in Table S1.

### Identification and evaluation of housekeeping genes

To calculate the skewness score, we computed the percentage of cells within every cell type expressing each gene present in our scRNA-Seq data. We then used baseR’s skewness function to score the skewness of every gene with respect to their percent of cells expressed within each cell type: a negative value (left skew) indicating expression in the majority of cells and cell types and a positive value (right skew) indicating expression in the minority of cells and cell types. To compute the Gini coefficient for every gene across cell types, we used the ineq function from the ineq package on the scaled TPM gene by cell type matrix. To perform gene ontology enrichment analysis, we used Wormbase’s gene set enrichment analysis tool with the default q-value threshold of 0.1. To measure the enrichment of essential genes in our various housekeeping genes list, we first downloaded the list of genes annotated as “embryonic lethal”, “larval lethal” and “adult lethal” from Wormbase. We combined these lists into a final list of essential genes made up of 3275 genes. We then used Fisher’s exact test to determine the extent of enrichment of essential genes in our housekeeping genes lists.

### Inferring transcriptional regulators underlying cellular identity

To infer the potential role of TFs in mediating cell-type specific gene expression, we correlated transcription factor binding patterns obtained from ChIP-Seq analysis with the gene expression profile of each cell type. We first collected all the available ChIP-Seq data from the modENCODE/modERN projects ^20–22^. All ChIP-Seq data are currently available from the ENCODE Data Coordination Center (DCC). We included in the analysis ChIP-Seq data performed in any post-embryonic stage (276 TFs) and ChIP-Seq data at embryonic stages (87 TFs) if the TF had not been tested post-embryonically. The ChIP-Seq peaks were then clustered along the genome by sorting the peaks by the apex base position of the peak. The peaks were accumulated into clusters moving along the genome until a gap of 200 bases between peaks was encountered, at which point a new cluster was begun. This resulted in 56729 clusters, varying in size between 1 TF and hundreds of TFs. Clusters that contained more than 70 TFs were excluded from the analysis since these are considered HOT (high occupancy target) sites and are not likely to represent tissue specific binding events. Similarly, clusters containing a single TF were also excluded since they are likely enriched in spurious binding. The target genes of the peak clusters were assigned by proximity of the cluster to the transcription start site (TSS) of the nearby genes. If the average of the apex of the peaks in the cluster met two criteria, the cluster was assigned to the gene with the closest TSS. The first criterion was that the peak cluster must be within 2000 bases of the nearest gene TSS. The second criterion was that the distance to the next closest gene TSS must be at least 1.5 times the distance to the nearest gene TSS. The peaks in each experiment (TF/stage) were ranked by signal strength and normalized to a cumulative probability. We then used a matrix containing the normalized signal strength as values, the TF as columns and the target genes as rows as the predictor variable matrix input for a generalized linear model (glmnet in R). If the cluster had multiple peaks of a given TF or there were multiple clusters assigned to the same target with the same TF, the maximum signal strength for the TF was used in the predictor variable matrix. The response vector for the model was the aggregated gene by cell type matrix we generated from our scRNA-Seq data. We then ran a separate model for each cell type, generating a determined coefficient for each TF-cell type association. These coefficients were used to generate the heatmaps found in Fig. 3A, Fig. 3C, Fig. 3D, Fig. 3E and Table S3. Any negative coefficients were set to 0 in the heatmap and only TFs found to be expressed in the cell type were used in the model (no TF expression is represented in gray in the heatmap). TFs with no positive values greater than > 0.015 in any cell type were omitted from the heatmap. Finally, we performed a 20-fold cross validation to determine the mean square error for the cell type model. That number was appended to each cell type (Fig. 3A, Fig. 3C, Fig. 3D, Fig. 3E and Table S3) with a lower number indicating a higher confidence in the predictions of the model.

### Inferring cell-cell communication from the gene expression of ligands and receptors in cells

To study CCIs, we used a list of 245 ligand-receptor interactions of *C. elegans* ^33^. We employed *cell2cell* by using the pipeline cell2cell.analysis.SingleCellExperiment found in the *cell2cell* python package, which allows running a permutation analysis for computing the significance of the inferred communication scores for each combination of LR interaction and sender-receiver cell pairs, as previously introduced ^34^. To run this analysis, the expression level of each gene was aggregated at the cell-type level by computing the log1p(CPM) average expression within each cluster. Then, the communication score was computed as the geometric mean of the expression of the ligand in a sender cell type and the receptor in a receiver cell type.

To run *Tensor-cell2cell* to identify latent patterns of communication, only the communication scores with a P-value < 0.05 (indicating cell-type specific CCC) were used to build a 3D-communication tensor (Fig. S4A). This 3D tensor was decomposed by using *Tensor-cell2cell* into 11 factors, each factor representing a signature or module of CCC that summarizes a biological process involving specific cell types and ligand-receptor pairs.

## Acknowledgements

We thank Dr Kacy Gordon for her help with the cluster annotation, particularly with annotating the germline and somatic gonad. We thank Nella Solodukhina for helping manage the lab and preparing reagents. We thank the modERN consortium for allowing us to use their unpublished ChIP-seq data. This work was supported by grants from the NIH (DK087928 to EJO; R35 GM119850 to NEL; and U41HG007355 and R01GM072675 to RHW) and the W. M. Keck Foundation (to EJO). AG is supported by a dissertation-year fellowship from the Society of Fellows (UVA) and a Harrison Family Jefferson Fellowship from the Jefferson Scholars Foundation. EJO is supported by a Biomedical Scholars award from the Pew Charitable Trusts, an award from the Jeffress Trust. EA is supported by the Chilean Agencia Nacional de Investigación y Desarrollo (ANID) through its scholarship program DOCTORADO BECAS CHILE/2018 -72190270, the Fulbright Chile Commission, and the Siebel Scholars Foundation. NEL is supported by the Novo Nordisk Foundation (NNF20SA0066621). RHW was supported in part by the William Gates III Endowed Chair in Biomedical Sciences. Finally, for providing some of the *C. elegans* strains used in this study, we would like to acknowledge the CGC, which is funded by NIH Office of Research Infrastructure Programs (P40 OD010440).

## Author contributions

The study was conceived by AG and EJO and its planning and execution was carried out by AG, NEL, RW, and EJO. CH performed the scRNA-Seq experiment and AG, CH, RW, and EJO analyzed the scRNA-Seq data. AG performed the housekeeping gene analysis and AG and EJO interpreted the housekeeping genes results. LG performed the transcription factor analysis and AG, LG, RW, and EJO interpreted the results. EA performed the CCI analysis and AG, EA, NEL, and EJO interpreted the results. AG performed the RNAi experiment for the CCI analysis and EJO the chemotaxis controls. AG and EJO wrote the original draft of the manuscript and all authors contributed to the final version of manuscript.

## Additional information

Supplementary Information is available for this paper.

## Data and code availability

- A user-friendly interface for visualizing and exploring the scRNA-Seq data is available at: https://wormseq.org/
- All files including the cell metadata, gene metadata and expression matrices of all three replicates and the code used to generate all the analyses performed in this study are publicly available through Zenodo at: https://doi.org/10.5281/zenodo.7296547
- Raw data will be deposited on GEO (while under review, link available upon request).
- Any additional information required to reanalyze the data reported in this paper is available from the lead contact upon request.

**Figure S1.**
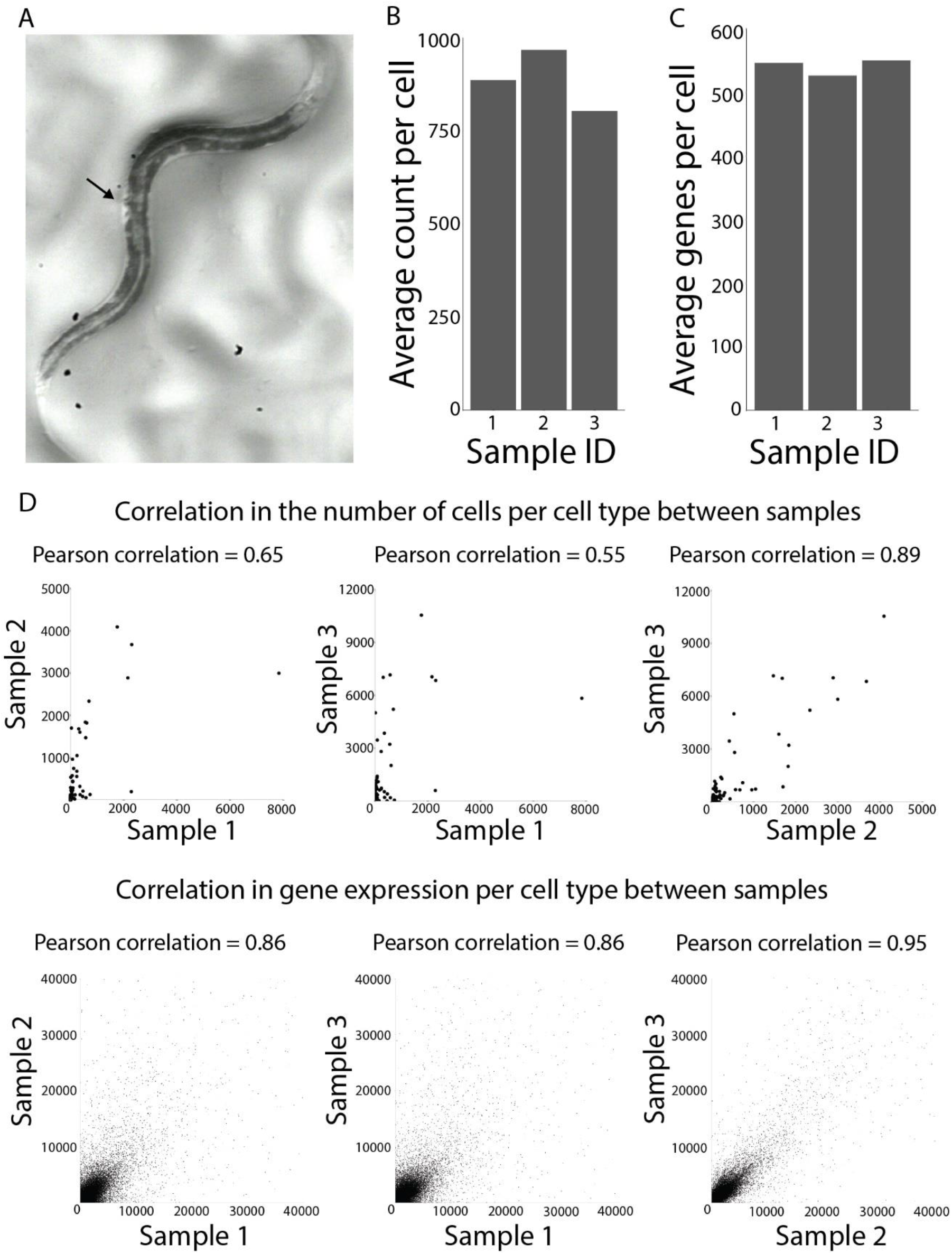
scRNA-Seq of young adult *C. elegans*. **(A)** Representative image of young adult *C. elegans*, which we identified by the characteristic shape of the vulva (arrow). **(B)** Average count per cell for each biological replicate. **(C)** Average gene per cell for each biological replicate. **(D)** Correlation in the number of cells per cell type between the three biological replicates. Each dot represents the number of cells in a particular cell type in two samples. **(E)** Correlation of the cell-type specific gene expression profiles between the three biological replicates. Each dot represents the levels of expression of a gene within a cell type (scaled TPM).

**Figure S2.**
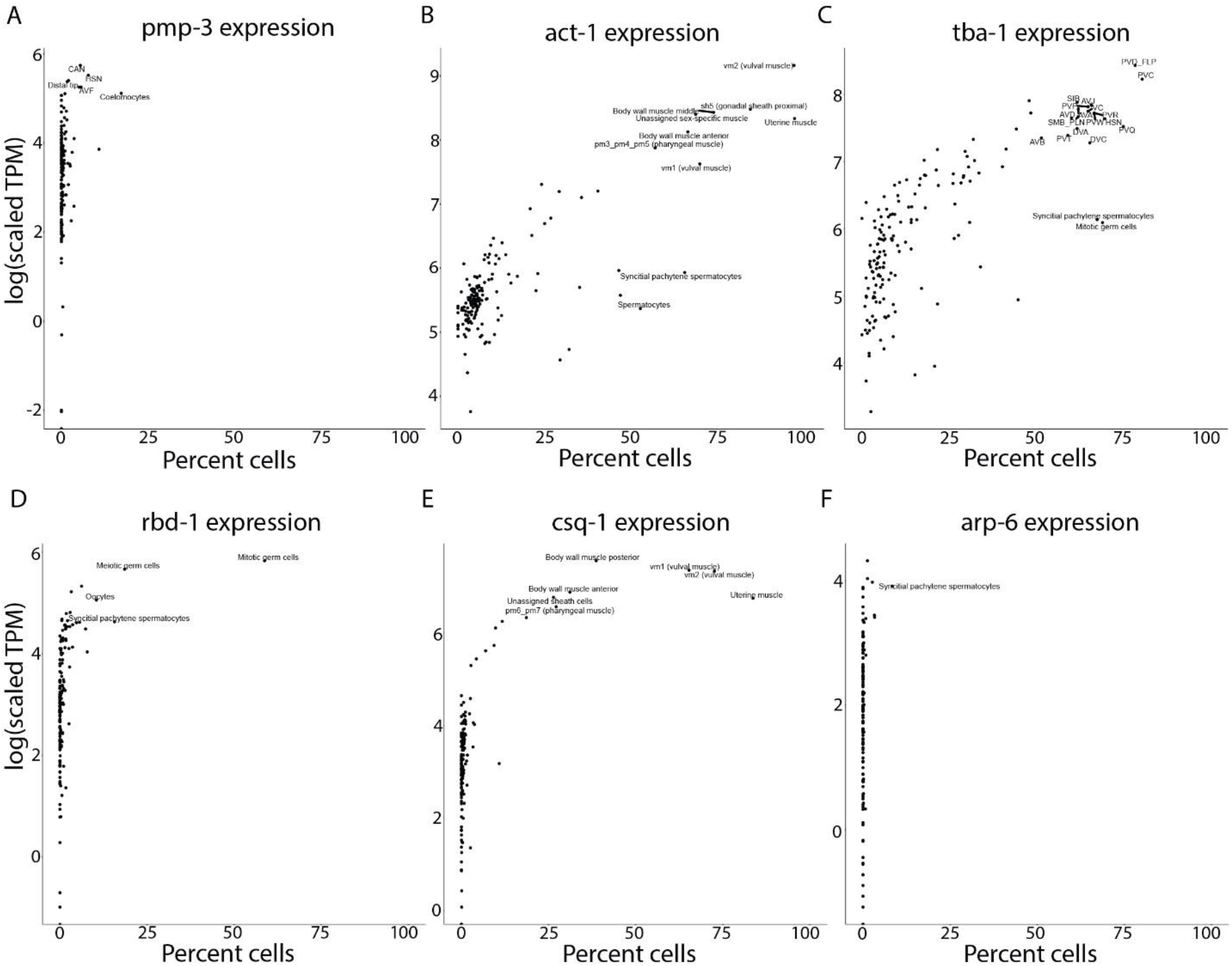
Curation of housekeeping genes. **(A-F)** Gene expression of traditional housekeeping genes with high Gini coefficient.

**Figure S3.**
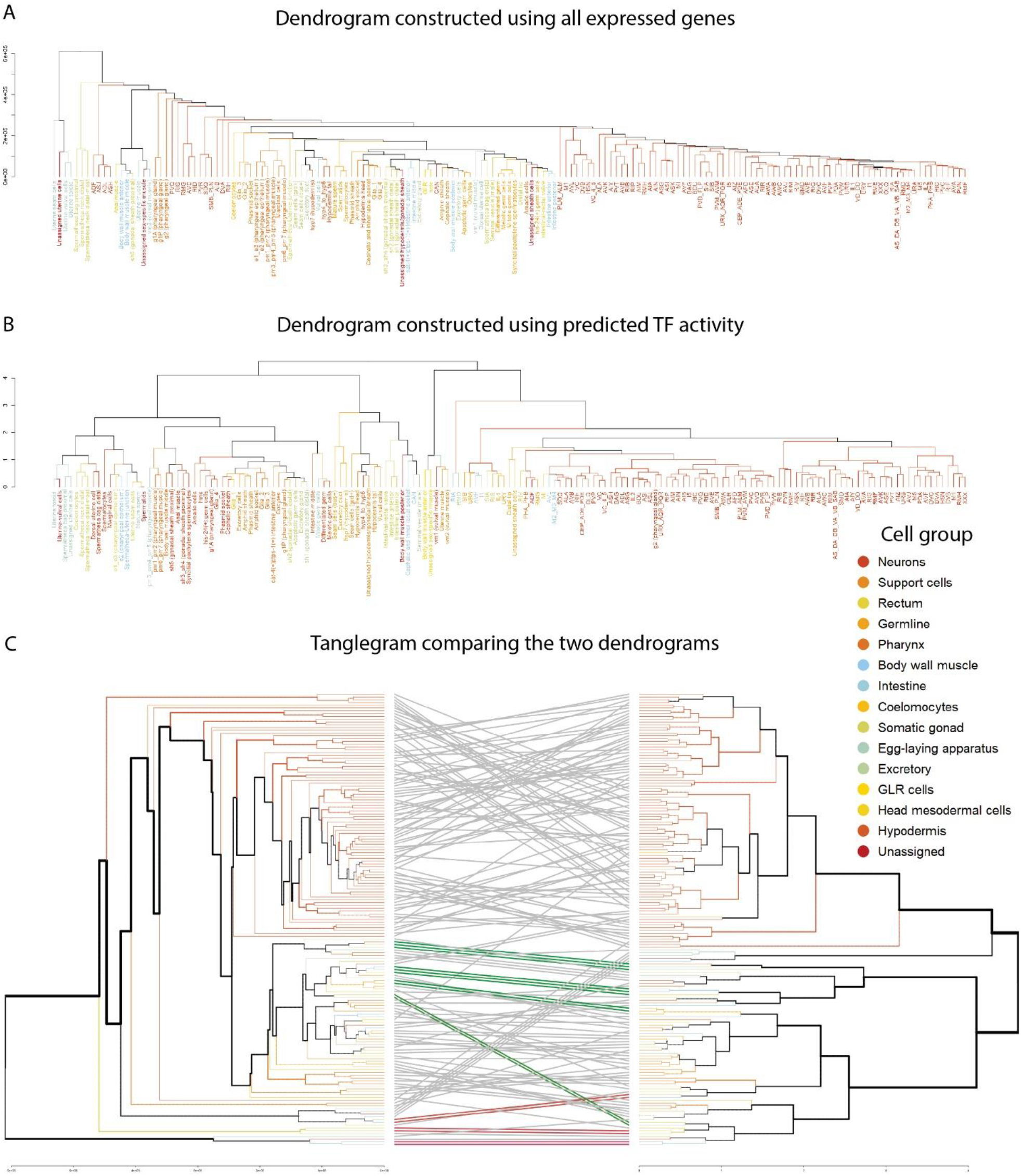
Comparison of the dendrogram constructed using all expressed genes and the dendrogram constructed using predicted TF activity alone. **(A)** Dendrogram constructed using all expressed genes. Colors represent cell group (legend on the right). **(B)** Dendrogram built using the predicted TF activity only. Color represents cell group (legend on the right). **(C)** Tanglegram showing the level of similarity between the TF and gene expression dendrograms. Non-shared nodes are depicted as dotted lines. Solid lines indicate leaves shared by both dendrograms. Colored lines indicate branches that are identical in both dendrograms.

**Fig. S4.**
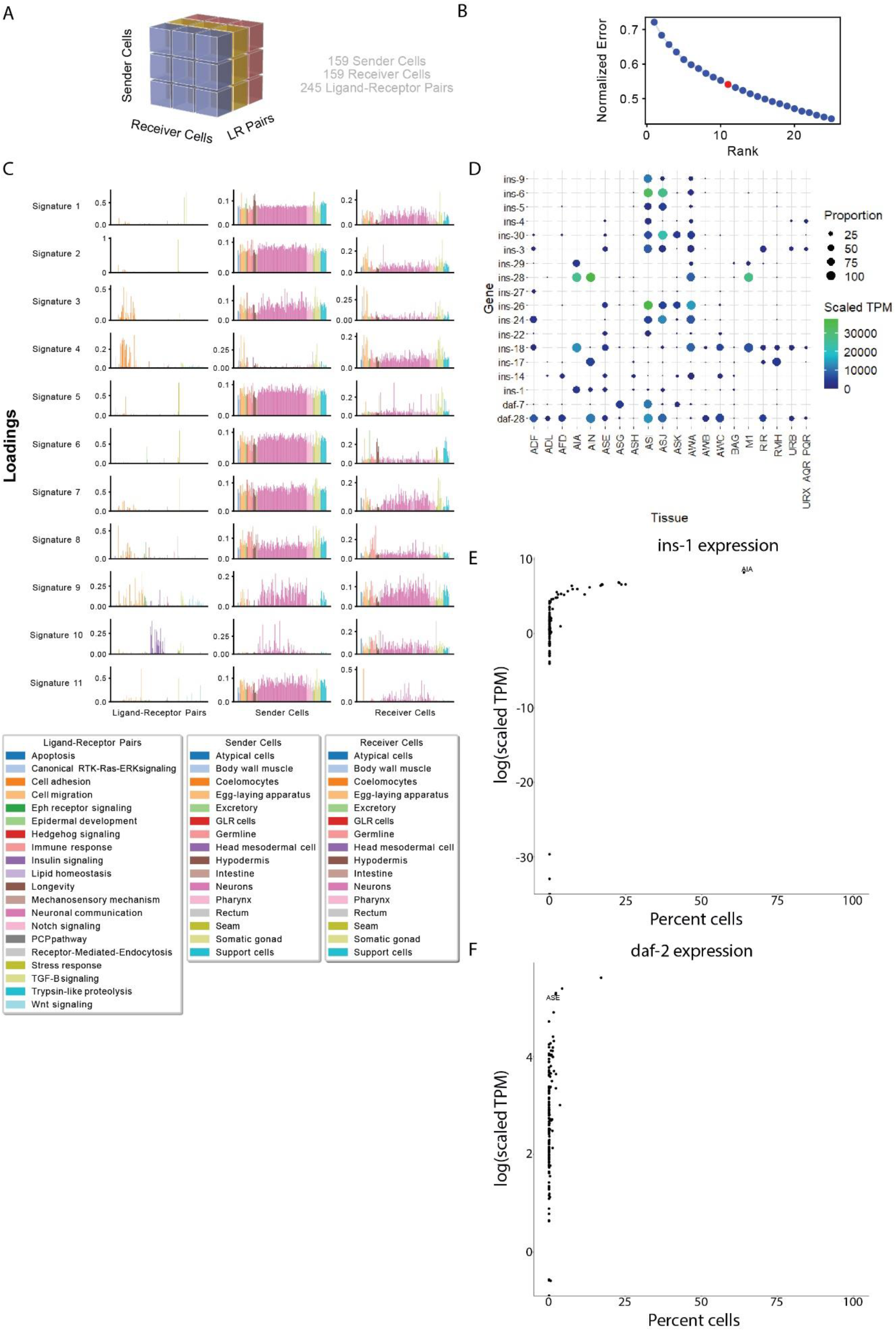
Identification of cell-type specific communication signatures using Tensor-cell2cell. **(A)** Graphical representation of Tensor analysis. **(B)** Elbow plot to determine the optimal number of factors/signatures. **(C)** Tensor-cell2cell analysis: The first column of graphs represents the importance of each ligand-receptor in each signature colored by ligand-receptor pair class, the second column of graphs represents the importance of each sender cell in each signature colored by tissue type, the third column of graphs represents the importance of each receiver cell in each signature colored by tissue type. Color legend of functional, sender-cell, and receiver-cell classes can be found below the graphs. **(D)** The distribution of insulin ligands important across the sender cells in signature 10. **(E)** Enrichment of *ins-1* expression in AIA neurons. **(F)** Enrichment of *daf-2* expression in ASE neurons.

